# Citrate synthase improves sepsis-induced lung injury by reconstructing the mitochondrial tricarboxylic acid cycle of macrophages

**DOI:** 10.1101/2024.10.16.618654

**Authors:** Jiaojiao Sun, Sihao Jin, Zhiqiang Wang

**Author notes:** Correspondence to: Zhiqiang Wang. Department of Cardiothoracic Surgery, Affiliated Hospital of Jiangnan University, No. 1000 Hefeng Road, Wuxi 214122, China. These authors contributed equally to this work (Jiaojiao Sun, Sihao Jin).

## Abstract

**Background:** The destruction of mitochondrial function during sepsis-induced acute lung injury (sepsis-ALI) can lead to tissue cell damage and organ dysfunction. Citrate synthase (CS) may maintain cellular energy metabolism by enhancing the mitochondrial tricarboxylic acid cycle (TCA cycle) in macrophages.

**Methods:** 76 healthy donors and 89 sepsis patients were included. The levels of CS were determined using ELISA. We established a cecal ligation and puncture (CLP) model of sepsis to evaluate the effects of CS on lung injury by lung macrophages-specific CS knockdown or CS inhibitors. Isolated mouse lung macrophages were stimulated with LPS to observe the impact of CS overexpression and knockdown on TCA cycle.

**Results:** In sepsis patients, CS was expressed at low levels and positively correlates with lung function parameters. In sepsis mice, knockdown CS or inhibiting its expression exacerbated lung injury and oxidative stress. In macrophages, inhibiting CS expression affected TCA cycle and worsened cell apoptosis, while overexpressing CS promoted TCA cycle, alleviating cell apoptosis, enhancing cellular energy production, and reducing oxidative stress levels. The supplementation of citric acid (a downstream metabolite of CS) helped alleviate mitochondrial damage and promotes the TCA cycle.

**Conclusions:** These results suggest that targeting CS may be a promising therapeutic approach for treating sepsis.

## Introduction

Acute lung injury (ALI) is a common complication of sepsis involving a cascade of inflammatory responses that can lead to significant damage to the lungs. This condition is typically caused by pulmonary infections or other factors that impair lung function (1, 2). The interplay of immune dysregulation, pathogen toxicity, and subsequent tissue damage contributes to the severity of septic lung injury (3). The management of septic lung injury necessitates a comprehensive approach, including addressing the underlying infection, supporting respiratory function, and managing systemic inflammatory responses. In this challenging clinical scenario, early detection and timely intervention are crucial for mitigating lung injury and improving patient outcomes.

The oxidative stress and lung macrophages damage occurring during the process of sepsis are closely associated with mitochondrial dysfunction (4, 5). Mitochondria serve as the powerhouse of cellular energy production and are also a primary source of oxidative stress. Impairment of mitochondrial function during sepsis can lead to disruptions in cellular energy metabolism, promoting inflammatory responses and cellular apoptosis (6, 7). Therefore, protecting mitochondrial function is crucial for mitigating tissue damage caused by sepsis. Understanding the role of mitochondria in sepsis and developing therapeutic strategies targeting mitochondria are therefore crucial for the management and treatment of sepsis.

Mitochondria are the powerhouses of the cell, and the TCA cycle is the key metabolic pathway within the mitochondria (8). Citrate synthase (CS) is the major rate-limiting enzyme of the TCA cycle, responsible for combining acetyl-CoA and oxaloacetate to form citrates (9). CS promotes the energy cycle of the TCA cycle and the electron transport chain (ETC), making it a quantitative marker for mitochondrial integrity, function, quality, and respiratory chain enzymes (10, 11). Reduced CS activity can impair aerobic ATP synthesis, decrease citrate levels, or reduce cytosolic acetyl-CoA, leading to a low-energy environment within the cell (12). Decreased CS levels result in reduced formation, release, and resynthesis of citrate, cytosolic acetyl-CoA, and the neurotransmitter Ach (13).

The relationship between CS and mitochondria encompasses energy production and involves several key biological processes within the cell. Therefore, further exploration of the interaction between CS and mitochondria helps deepen our understanding of cellular metabolic regulatory mechanisms. In this study, we aimed to investigate whether CS is involved in the pathogenesis of sepsis and explore its potential mechanisms. We found that CS has the potential to protect against sepsis-ALI and promote tissue repair. Additionally, citric acid, a downstream metabolite of CS, rescued mitochondrial dysfunction and alleviated sepsis-ALI.

## Materials and Methods

### Data processing and Identification of differential expression genes (DEGs)

The sepsis RNA expression data were downloaded from the Gene Expression Omnibus (http://www.ncbi.nlm.nih.gov/geo/). Data from GSE95233 and GSE208581 were used for differential expression analysis.

### Patients and Ethics

Serum samples from sepsis patients (*n* = 76) and healthy donors (*n* = 89) were collected at Jiangnan University Affiliated Hospital on June 1, 2023, and September 1, 2024 (Table S1). The clinical samples involved in this study were approved by the Ethics Committee of Jiangnan University Affiliated Hospital (Ethics Review No: LS2020038).

### Reagents and Antibodies

Octanoyl coenzyme A lithium (Octanoyl-CoA) (Cat# HY-134136A), Citric acid (Cat# HY-N1428), and Lipopolysaccharides (LPS) (Cat# H) were purchased from Med Chem Express (MCE, USA). The primary antibodies used were as follows: Citrate Synthase (#14309, rabbit monoclonal, Cell Signaling Technology, Danvers, MA, USA, 1:1000), Bcl2 (#15071, mouse monoclonal, Cell Signaling Technology, Danvers, MA, USA, 1:1000), BAX (#2772, rabbit monoclonal, Cell Signaling Technology, Danvers, MA, USA, 1:1000), VDAC1 (#4866, rabbit monoclonal, Cell Signaling Technology, Danvers, MA, USA, 1:1000), and β-actin (#66009-1-Ig, mouse monoclonal, proteintech, 1:5000), COX I (#0807-3, rabbit polyclonal, HUABIO, 1:1000), and TFAM (#ab176558, rabbit monoclonal, Abcam, Cambridge, MA, USA, 1:1000).

### Animal Experimental Design

The animal experiments were approved by the Animal Ethics Committee of Jiangsu Institute of Schistosomiasis Control (approval number: IACUC-JIPD-2019043). 6-8-week-old WT mice (C57BL/6 background) were kept under specific pathogen-free (SPF) conditions. The WT mice were randomly divided into four groups (n = 6): Sham group (Sham), Octanoyl-CoA group (Octanoyl-CoA), CLP group (CLP), and Octanoyl-CoA intervention group (CLP+ Octanoyl-CoA).

The CLP-operated sepsis mouse model was prepared as previously described (14). Briefly, a 2 cm midline incision was made to expose the cecum and adjoining intestine. The cecum was then ligated tightly using a 3.0-silk suture 5.0 mm from the cecal tip, punctured with a 22-gauge needle, and then gently squeezed to extrude feces from the perforation site. The cecum was then returned to the peritoneal cavity, and the laparotomy site was sutured using 4.0-silk. For sham operations, the cecum of animals was surgically exposed, but not ligated or punctured, and then returned to the abdominal cavity. The mice were sacrificed, and bronchoalveolar lavage fluid, blood samples, and lung tissue specimens were collected.

### The Accumulative Survival Rates

Male C57BL/6 mice were randomly divided into 3 groups (each n = 10): Sham, CLP, and CLP + Octanoyl-CoA. The survival time was recorded at 24 h intervals, and all mice that survived were euthanized after 96 h.

### Blood Gas Analysis

Arterial blood samples were collected using heparinized syringes from the femoral artery of mice. Blood gas analysis was performed using a blood gas analyzer (the ABL90 FLEX analyzer) capable of measuring parameters including pH, partial pressure of oxygen (PaO_2_), and partial pressure of carbon dioxide (PaCO_2_).

### Acquisition and analysis of BALF

After fixation, the mice were intubated, the right lung tissues were rinsed three times with 0.6 mL sterile normal saline, and the collected BALF solution was centrifuged at 1000 g for 10 min at 4 °C.

### Lung histological assay and wet-to-dry ratio analysis

Mice lung specimens were fixed in 4% paraformaldehyde for 48 h and subsequently embedded in paraffin using ASP200S and EG1150H (Leica). At the time of death, the left lungs of mice were taken and weighed to determine the wet weight, then placed in an oven at 80°C for 24 h and reweighed to determine the dry weight for calculation of the wet/dry weight ratio.

### Immunohistochemistry Staining

Sections of lung tissues, 4 µm thick, were deparaffinized, rehydrated, and subjected to antigen repair in 10 mM sodium citrate buffer heated to 95 °C for 30 min. Non-specific binding was blocked with normal goat serum for 20 min at 37 °C.

### Biochemical Indexes Analysis

The concentration of MDA (Cat. No: A003-1-2), MPO (Cat. No: A044-1-1), GSH (Cat. No: A006-2-1) and SOD (Cat. No: A001-3-2) were determined using commercial kits provided by Nanjing Jiancheng Bioengineering Institute (Nanjing, China) with an UV-VIS spectrophotometer (Thermo, Waltham, MA, USA) according to the manufacturer’s instructions.

### Mitochondrial Metabolomics Analysis

The lung mitochondria were used for metabolomics analysis. The midbrain mitochondria were separated by using the mitochondrial extraction kit. The metabolites were extracted with 1 mL of precooled methanol and then crushed by ultrasound for 5 min. Thereafter, the precipitate was centrifuged, and the supernatant was collected, which was dried into powder by a centrifugal concentrator. The powder was dissolved in a 600-mul phosphate buffer and centrifuged, and a 550-mul solution was put into an NMR tube for determination.

### Pulmonary Macrophages Isolation and Treatment

Pulmonary macrophages were isolated from mouse lungs according to previously described methods. Briefly, mouse lung tissues were homogenized, and single-cell suspensions were prepared. Macrophages were separated by density gradient centrifugation with Histopaque-1083 (Sigma-Aldrich, St Louis, MO) and plated at a concentration of 6 × 10^6^ cells/mL in Dulbecco’s Modified Eagle Medium (DMEM) containing 20% fetal calf serum (FCS). After 3 h, 95% of adherent cells were macrophages, as confirmed by flow cytometry.

### Enzyme-Linked Immunosorbent Assay

The contents of CS in mouse serum and cells were measured with ELISA kits (Cat. No. MAK057; Sigma-Aldrich, Merck KGaA).

### Adenosine Triphosphate Content (ATP) and cell counting Kit-8 (CCK-8) assay

ATP contents were quantified using an Enhanced ATP Assay Kit (#S0026, Beyotime, Nanjing, China) following the manufacturer’s instructions. Briefly, cells were seeded in 6-well plates at a density of 2 × 10^5^/cells/well and cultured overnight. Subsequently, the cells were exposed to LPS. Lastly, cell lysis was performed using a lysis reagent followed by centrifugation at 12,000 × g for 5 min at 4 °C. The supernatant was then collected to measure ATP levels. The cell viability of macrophages was detected via a CCK-8 assay kit (#C0037, Beyotime, Nanjing, China). 10 µl CCK-8 reagent was added to each well of the 96-well plate and incubated at 37 °C for 1.5 - 2 h. The absorbance at 450 nm was detected by a Microplate Reader.

### CS Activity Assay

1 × 10^6^ cultured cells were collected and homogenized to extract mitochondrial. The protein concentration was measured by Pierce™ BCA Protein Assay Kit and the CS activity was measured by citrate synthase assay kit pro-caspase3 (#ab119692, Abcam, Cambridge, MA, USA) following the protocol recommended by the manufacturer.

### Measurement of ETC complexes Ⅰ - Ⅴ Activity

The concentration of complex Ⅰ (Cat. No: A089-1-1), complex Ⅱ (Cat. No: A089-2-1), complex Ⅲ (Cat. No: A089-3-1), complex Ⅳ (Cat. No: A089-4-1), and complex Ⅴ (Cat. No: A089-5-1) were determined using commercial kits provided by Nanjing Jiancheng Bioengineering Institute (Nanjing, China) with an UV-VIS spectrophotometer (Thermo, Waltham, MA, USA) according to the manufacturer’s instructions.

### Measurement of PDH, α-KGDH, NAD^+^/NADH, and ATP/ADP Activity

The concentration of PDH (Cat. No: H262-1-2), α-KGDH (Cat. No: H465), NAD^+^/NADH (Cat. No: A114-1-1), and ATP/ADP (Cat. No: A095-1-1) were determined using commercial kits provided by Nanjing Jiancheng Bioengineering Institute (Nanjing, China) with an UV-VIS spectrophotometer (Thermo, Waltham, MA, USA) according to the manufacturer’s instructions.

### Immunofluorescence Staining

Cells were fixed with 4% paraformaldehyde at room temperature for 15 min. Then, they were permeabilized with 0.1% Triton X-100 for 20 min. Subsequently, the fixed cells were washed with PBS and blocked with 5% goat serum at room temperature for 30 min. The corresponding fluorescent secondary antibodies were incubated at room temperature for 2 h. After washing with PBS, the cells were sealed using DAPI reagent (#P0131, Beyotime, Shanghai, China) and imaged using a Zeiss LSM 880 laser confocal fluorescence microscope (Carl Zeiss, Oberkochen, Germany).

### MitoSOX Red and Mito Tracker Red Detection

To detection of mitochondrial ROS, macrophages were stained with 5 µM of MitoSOX Red Mitochondrial Superoxide Indicator (#S0061S, Beyotime, Shanghai, China) for 10 min. After staining, macrophages were washed with PBS, and then macrophages were counterstained with DAPI for 10 min. The stained macrophages were observed and photographed on a confocal microscope. Macrophages were treated with 100 nM Mito tracker Red CMXRos (#C1035, Beyotime, Shanghai, China) for 30 minutes at 37 ° C, followed by washing and incubation in fresh medium for an additional 30 minutes. Stained macrophages were observed using a fluorescence microscope.

### Transmission Electron Microscopy (TEM)

Macrophages were first fixed with 2.5% glutaraldehyde in phosphate buffer for > 4 h, followed by post-fixation with 1% OsO4 in phosphate buffer for over 2 h. Subsequently, the samples were stained with uranyl acetate for 5 min or alkaline lead citrate for 10 min. Images were collected and analyzed using a Hitachi transmission electron microscope (Hitachi, Model H-7650, Tokyo, Japan).

### Flow Cytometry

Macrophages were inoculated into 6-well plates (2 × 10^5^/well). After LPS stimulation, the macrophages were collected and reacted with 5 µl Annexin V-FITC (#C1062S, Beyotime, Shanghai, China) and 5 µl propyl iodide (PI) at room temperature and in the dark for 20 min. Apoptosis was detected by flow cytometry (BD Biosciences, San Jose, CA, USA).

### Measurement of Extracellular Acidification Rate

Extracellular acidification rate (ECAR) was analyzed using the pH-sensitive BBcellProbe P61 fluorescent probe supplied by the ECAR assay kit (BB-48311, BestBio). Cells were seeded in a 96-well clear-flat-bottom black microplate and incubated for 2 d. The BBcellProbe P61 probe was then added to the cells. The plate was read on an EPOCH2 microplate reader (BioTek) in kinetic mode at 37 °C for 2 h (1 read per 3 min, Ex 488/Em 580). ECAR was calculated from the slope of the kinetic curve according to the manufacturer’s instructions.

### Measurement of Oxygen Consumption Rate

Oxygen consumption rate (OCR) was measured with the Oxygen Consumption Rate kit (600800, Cayman Chemical). Macrophages were seeded in a 96-well clear-flat-bottom black microplate and incubated for 2 d. After adding the Phosphorescent Oxygen probe to the wells, 100 μl mineral oil was added to each well on top of the medium. The plate was read on an EPOCH2 microplate reader (BioTek) in kinetic mode at 37 °C for 2 h (1 read per 3 min, Ex 380/Em 650). Basal OCR levels of the macrophages were calculated from the slope of the kinetic curve following the manufacturer’s instructions.

### CS Short Interfering RNA Transfection

CS-specific short interfering RNA (siRNA) molecules were chemically synthesized by Shanghai RiBobio Company (CS siRNA: ATTCCTCTGGCCTTGG). Transfection was performed by utilizing Lipofectamine RNAiMAX reagent (Life Technologies, CA, USA) as the manufacturer’s protocol. Macrophages were allowed to recover in a fresh growth medium of 48-72 h to permit CS silencing.

### Plasmid construction and transfection

The CS-OE plasmid pcDNA3.1(+)-CS-3 × FLAG-P2A-EGFP (forward primer [CMV-F], CTTGTCCTTAACGCGTG; reverse primer [EGFP-SEQR], GGTTCGTCTGAATAACAGCTT) and the control vector H2713 pcDNA3.1(+) - 3 × FLAG-P2A-EGFP were designed and constructed by OBiO Technology. HEK-293T cells were seeded at a density of 5 × 10^5^ cells per well in six-well plates and transfected with 2 μg DNA per well mixed with 5 μl Liposomal Transfection Reagent (Yeasen; Cat. No: 40802ES02) according to the manufacturer’s protocol.

### Quantitative Real-Time PCR

The total RNA from cells or tissues was extracted by Trizol reagent (Vazyme, Nanjing, China), and 1 µg extracted RNA was reverse transcribed into cDNA using the RNA PCR Kit (Yeasen, Shanghai, China). Quantitative PCR was carried out using SYBR^®^ Green RT PCR Master Mix (Yeasen, Shanghai, China) and a Light Cycler 480-II Real-Time PCR system (Roche, Basel, Switzerland). Relative RNA expression levels were calculated by applying the 2^−ΔΔCt^ method, and the Β-ACTIN gene was used as an endogenous control gene for normalizing the expression of target genes. The primer sequences are shown in Table S2. Genomic DNA was isolated using Qiagen Genomic-tip 20/G and Qiagen DNA Buffer Set (Qiagen, Gaithersburg, MD) per the manufacturer’s instruction. Eluted DNA was incubated with isopropanol overnight at -80 C and centrifuged 12,000g for 60 min. DNA was washed with 70% ethanol and dissolved in TE buffer. PCR was performed using Ex-taq (Clonetech, Mountain View, CA).

### Western blot

The samples were homogenized in ice-cold RIPA buffer with a protease inhibitor mixture, and the supernatant was collected after centrifugation (12,000 rpm, 10 min, and 4 °C). Protein concentration in the supernatant was determined using a BCA protein kit and subsequently denatured at 100 °C for 10 min. The protein was separated on a 12.5% SDS-PAGE gel and transferred onto a nitrocellulose filter. The filter was then blocked with 5% skimmed milk powder for 2 h before being incubated overnight at 4 °C with the primary antibody. After washing the membrane with TBST, the filter was incubated with the corresponding secondary antibody at room temperature for 2 h. The protein level was detected using a Bio-Rad imaging system equipped with a chemiluminescent substrate (Tanon, Shanghai, China).

### Statistical analysis

The data were presented as means ± standard deviation of at least three independent experiments. Group comparisons were conducted using analysis of variance (ANOVA). Categorical variables in clinical data were evaluated using the chi-squared test. Correlations were determined through linear regression analysis. Cumulative survival rates were assessed using the log-rank test. A *p* value < 0.05 was considered statistically significant for all tests conducted. Data analysis and image processing were performed using Prism v9.0 (GraphPad) and SPSS v20 (IBM).

## Results

### 1 CS levels were decreased in sepsis patients by bioinformatics analysis

Through bioinformatics analysis of data from healthy donors and sepsis patients in GSE95233 and GSE208581, differential gene selection was performed on the two datasets under the criteria of adjusted p < 0.05 and absolute log2 fold change (FC) > 0. The table shows the basic information of GSE95233 and GSE208581 (Figure 1A). Kyoto Encyclopedia of Genes and Genomes (KEGG) enrichment analysis revealed the significant role of the tricarboxylic acid cycle (TCA cycle) in sepsis (Figure 1B-1C). The intersection of DEGs from the two datasets was calculated, resulting in 18 overlapping DEGs (Figure 1D-1E).

**Figure 1.**
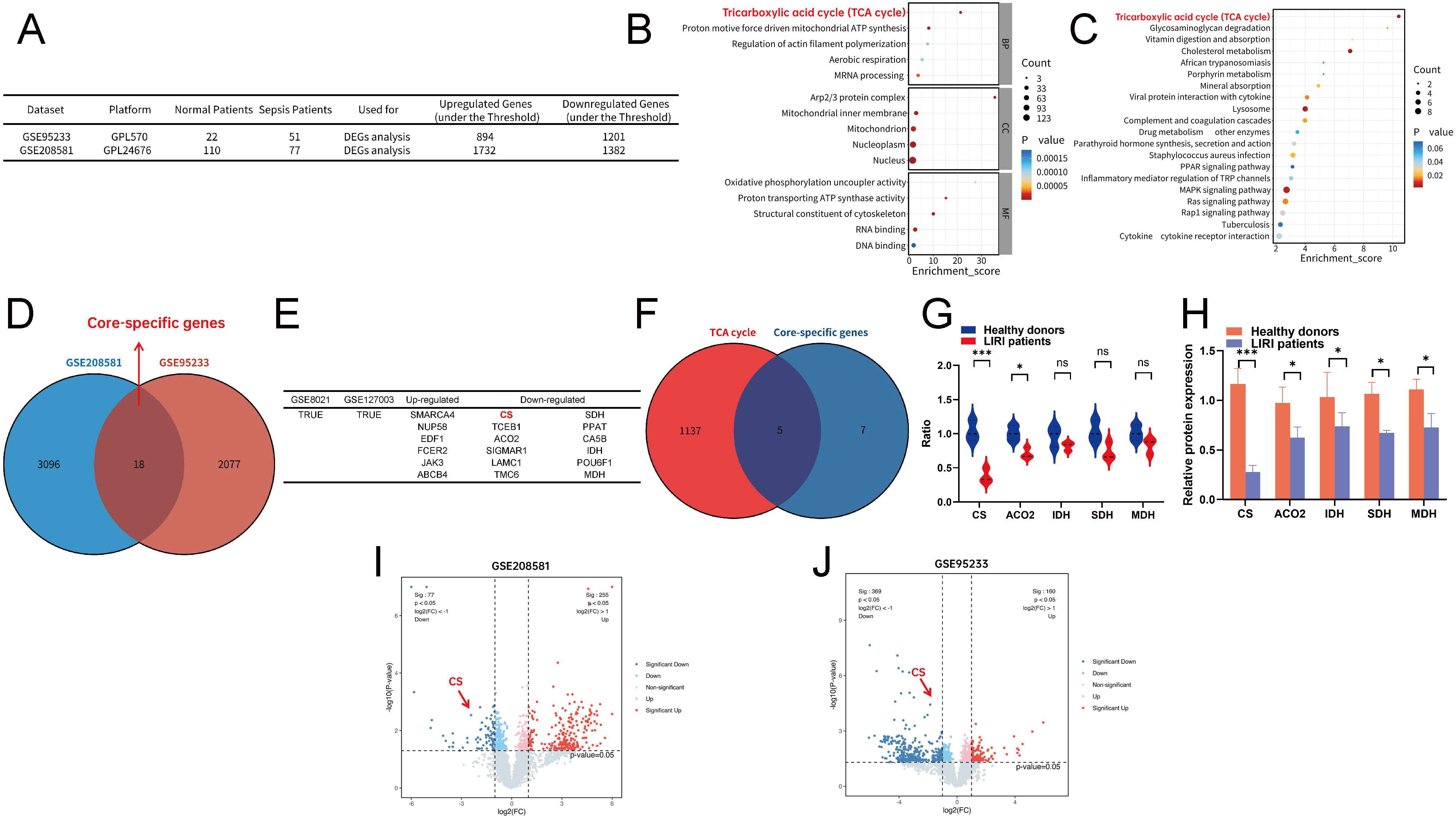
CS levels were decreased in sepsis patients by bioinformatics analysis. (A) The overview of the dataset used for this study. (B) KEGG analysis of GSE95233. (C) KEGG analysis of GSE208581. (D) Venn plot of overlapping genes in two datasets. (E) The overview of the 18 overlapping DEGs. (F) Venn plot of overlapping genes in the TCA cycle and the 18 overlapping DEGs. (G) The expression of CS, ACO2, IDH, SDH, and MDH. (H) Quantification of serum CS, ACO2, IDH, SDH, and MDH levels by ELISA in healthy donors (*n* = 76) and sepsis patients (*n* = 89). (I) Volcano map of DEGs between healthy donors and sepsis patients in GSE95233 dataset. (J) Volcano map of DEGs between healthy donors and sepsis patients in GSE208581 dataset. Group comparisons were conducted using analysis of variance (ANOVA). Experiments were repeated at least three times. \**p* < 0.05 and *** *p* < 0.001. ns means no significant difference.

By searching all genes related to the TCA cycle from the GeneCards website and conducting a set analysis with the 18 overlapping DEGs, five overlapping DEGs were discovered, namely CS, ACO2, IDH, SDH, and MDH (Figure 1F). Additionally, CS was found to be significantly decreased in sepsis (Figure 1G). ELISA testing of serum from sepsis patients revealed the most prominent decrease in CS (Figure 1H). Volcano plots demonstrate the decrease of CS in GSE95233 and GSE208581 (Figure 1I-1J).

### 2 CS levels were decreased in sepsis patients

To determine whether CS levels changed in sepsis patients, serum samples were collected from 76 healthy donors and 89 sepsis patients, and the relevant characteristics are shown in Supplementary Table 1. ELISA results showed that the levels of CS in sepsis patients were significantly lower than those in healthy donors (Figure 2A), and the levels of CS activity were also measured, and the results showed that the levels of CS activity in sepsis patients were all significantly lower than those in healthy donors (Figure 2B).

**Figure 2.**
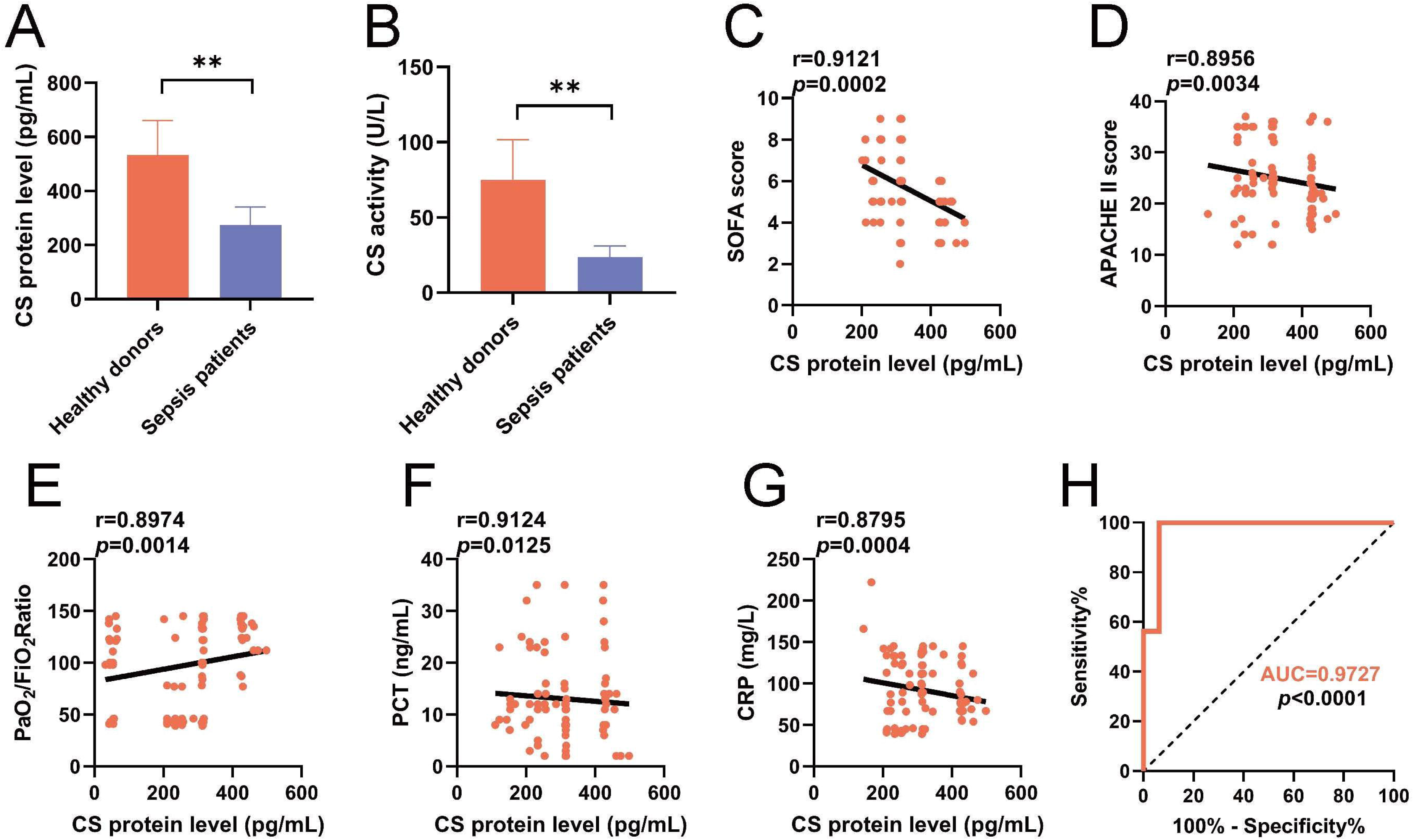
CS levels were decreased in sepsis patients. (A) Quantification of serum CS levels by ELISA in sepsis patients and healthy donors. (B) Quantification of serum CS activity in sepsis patients and healthy donors. (C-D) Serum CS levels were negatively correlated with higher SOFA /APACHE II scores in sepsis patients. (E) Quantitative analysis for the correlation between PaO_2_/FiO_2_ and CS levels in sepsis patients. (F) Quantitative analysis for the correlation between PCT and CS levels in sepsis patients. (G) Quantitative analysis for the correlation between CRP and CS levels in sepsis patients. (H) ROC curves to validate the diagnostic value of CS between sepsis patients and healthy donors. Group comparisons were conducted using analysis of variance (ANOVA) (healthy donors (n = 76) and sepsis patients (*n* = 89)). Experiments were repeated at least three times. \*\**p* < 0.01.

Moreover, a negative correlation was identified between serum CS levels and sepsis severity, as indicated by both the Sequential Organ Failure Assessment (SOFA) score (r = 0.9121, *p* = 0.0002; Figure 1C) and Acute Physiology and Chronic Health Evaluation II (APACHE II) score (r = 0.8956, *p* = 0.0034; Figure D). Moreover, we analyzed the correlation between CS levels and PaO_2_/FiO_2_, procalcitonin (PCT), and C-reactive protein (CRP) levels in sepsis patients. PaO_2_/FiO_2_ (r = 0.8974, *p* = 0.0014; Figure 2E) was positively correlated with CS expression. PCT (r = 0.9124, *p* = 0.0125; Figure 2F) and CRP (r = 0.8795, *p* = 0.0004; Figure 2G) were negatively correlated with CS expression. The receiver operating characteristic (ROC) curve was used to analyze the diagnostic value of serum CS level for sepsis. The results showed that the area under the curve (AUC) for comparing sepsis patients’ group was 0.9727 (Figure 2H). The above results indicate that CS is under-expressed in sepsis patients, and the expression level of CS is correlated with the lung function of the patients. Lower expression of CS suggests poorer lung function in the patients.

### 3 CS levels decreased in sepsis mice and pulmonary macrophages

Meanwhile, C57BL/6 mice were used to observe the changes in CS levels in sepsis mice. It was found that both mRNA and protein levels of CS were decreased in sepsis mice (Figure 3A-3B). Tissue immunofluorescence also indicated a decrease in CS protein levels (Figure 3C). Additionally, it was observed that CS activity was also reduced in sepsis mice (Figure 3D). To determine if CS levels would change in cells as well, we examined the changes in CS levels in pulmonary macrophages after LPS stimulation. It was found that both mRNA and protein levels of CS decreased in pulmonary macrophages after LPS stimulation (Figure 3E-3F). CS activity was also decreased in pulmonary macrophages after LPS stimulation (Figure 3G). Immunofluorescence also showed a decrease in CS protein levels (Figure 3H). The above results indicate that CS expression was low both in sepsis mice and pulmonary macrophages.

**Figure 3.**
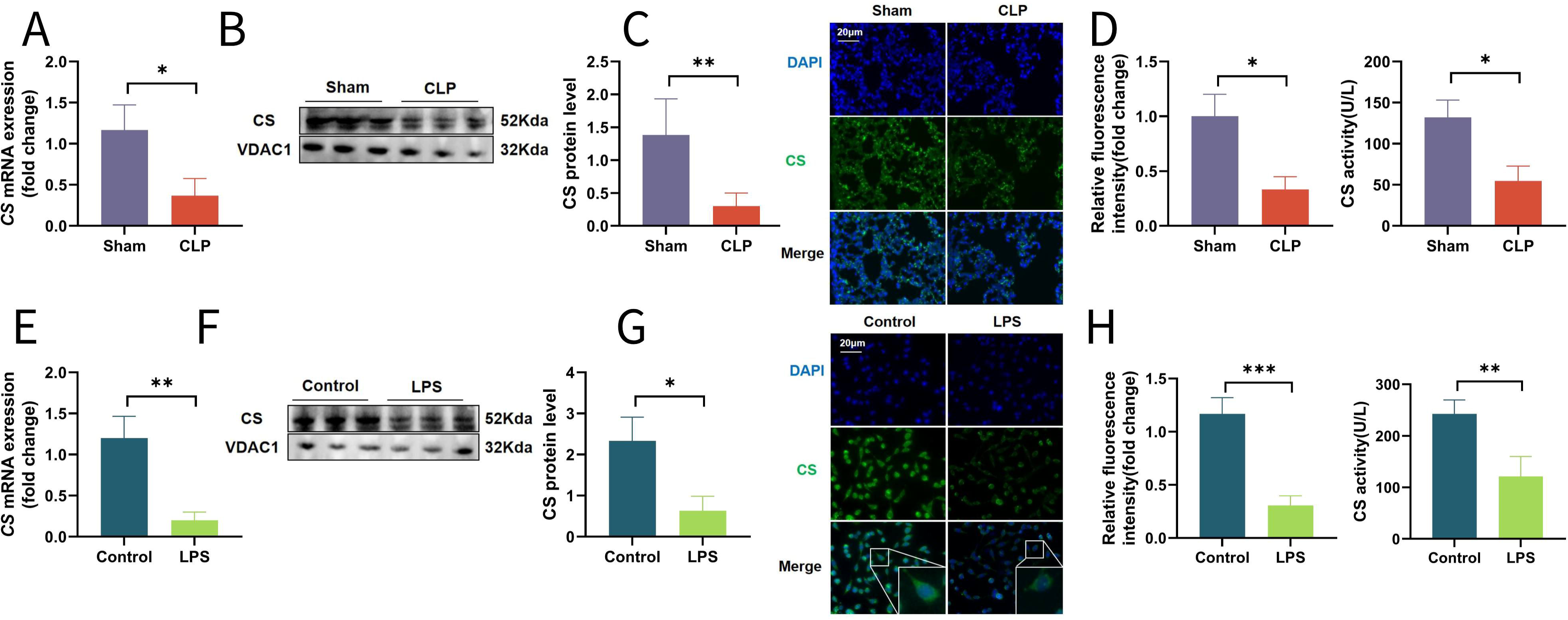
CS levels decreased in sepsis mice and pulmonary macrophages. (A) The mRNA levels of CS in lung tissues were detected by quantitative real-time PCR (*n* = 6). (B) The protein levels of CS in mice lung tissues were detected by western blot, and the ratio of CS was semi-quantitatively analyzed by ImageJ software (*n* = 6). (C) CS immunohistochemistry and semi-quantitative analysis of CS immunohistochemistry (magnification, ×200, *n* = 6). (D) Quantification of serum CS activity in lung tissues (*n* = 6). (E) The mRNA levels of CS in pulmonary macrophages were detected by quantitative real-time PCR (*n* = 3). (F) The protein levels of CS in pulmonary macrophages were detected by western blot, and the ratio of CS was semi-quantitatively analyzed by ImageJ software (*n* = 3). (G) Quantification of serum CS activity in pulmonary macrophages (*n* = 3). (H) Pulmonary macrophages were stained with the CS (green) and DAPI (blue) for immunofluorescence analysis, scale bar = 20 µm (*n* = 3). Group comparisons were conducted using analysis of variance (ANOVA). Experiments were repeated at least three times. \**p* < 0.05, \*\**p* < 0.01, and \*\*\**p* < 0.001.

### 4 CS knockdown aggravated lung injury in sepsis mice

We then sought to specifically inhibit CS in lung macrophages by stereotaxic injection of adeno-associated virus (AAV)-lungX-glial fibrillary acidic protein (GFAP)-shCS targeted to lung macrophages (Figure 4A). CS short hairpin RNA significantly reduced CS mRNA and protein expression in lung macrophages compared with the Sham group (Figure 4B-4C). Blood gas analysis demonstrated a decrease in PaO_2_ and an increase in PaCO_2_ after CS knockdown (Figure 4D-4E). Additionally, histopathological examination revealed thickening of the alveolar walls in mice after CS knockdown (Figure 4F). Furthermore, there was an increase in lung injury scores (Figure 4G) and lung wet/dry ratio (Figure 4H) following CS knockdown. Moreover, the total cell counts and protein concentration levels in BALF significantly increased after CS knockdown (Figure 4I-4J).

**Figure 4.**
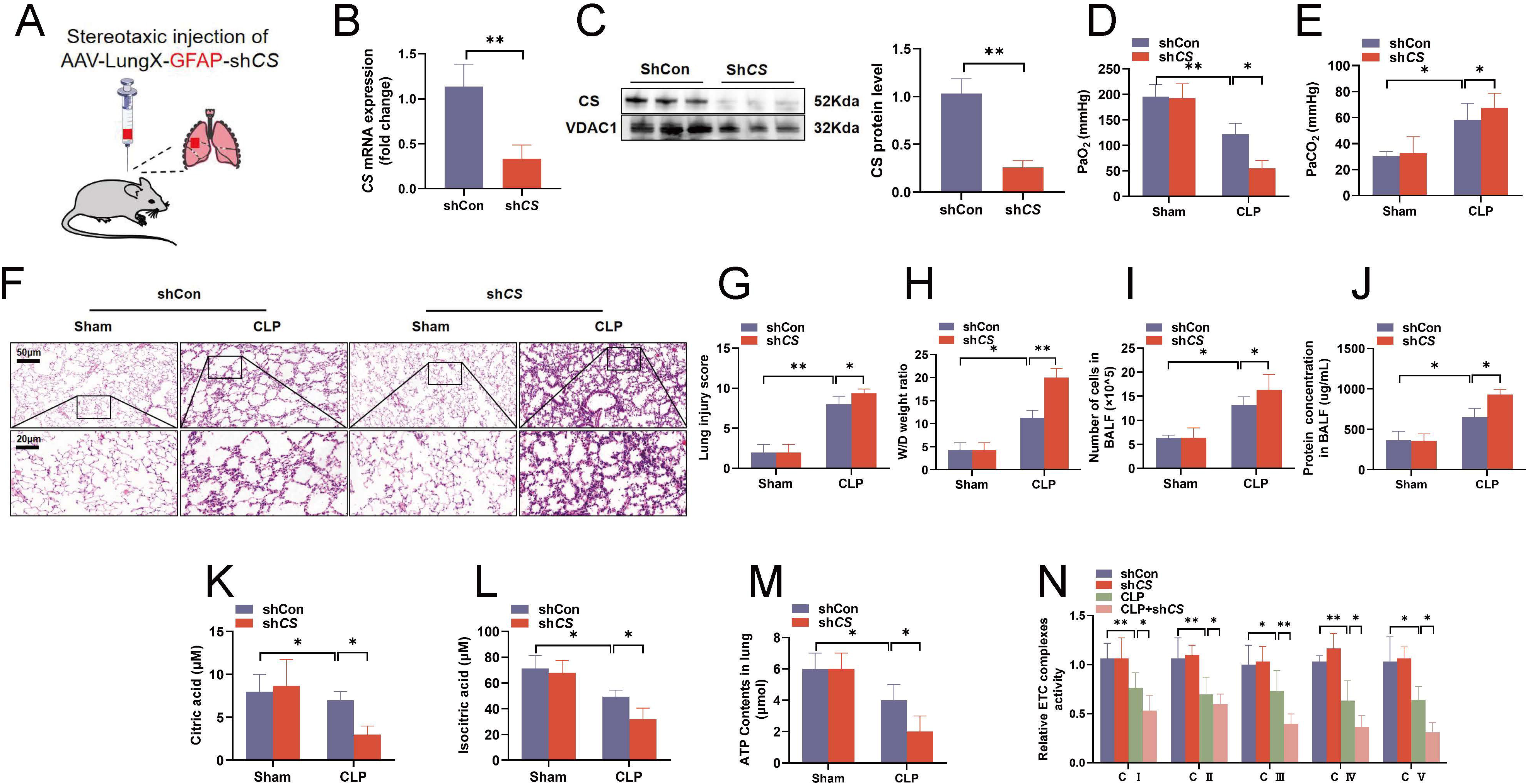
CS knockdown aggravated lung injury in sepsis mice. (A) Schematic illustration showing the stereotaxic injection of adeno-associated virus (AAV)-lungX-glial fibrillary acidic protein (GFAP)-shCS targeted to lung macrophages of C57BL/6J male mice. (B) The mRNA levels of CS in lung tissues were detected by quantitative real-time PCR. (C) The protein levels of CS in mice lung tissues were detected by western blot, and the ratio of CS was semi-quantitatively analyzed by ImageJ software. (D-E) Blood gas analysis of sepsis mice. (F) Hematoxylin and eosin staining were performed to evaluate the lung histopathological changes (magnification, × 200). (G) Mouse lung injury score was calculated. (H) The wet-to-dry ratio of lung tissues. (I) The total number of cells in BALF. (J) The total protein concentrations of BALF. (K-L) The levels of citric acid and isocitric acid in lung tissue. (M) The contents of ATP in lung tissues. (N) The levels of ETC complexes in lung tissues. Group comparisons were conducted using analysis of variance (ANOVA). Experiments were repeated at least three times (*n* = 6). \**p* < 0.05 and \*\**p* < 0.01.

In sepsis mice, we also detected changes in the levels of citric acid and isocitric acid, which are indicators of TCA cycle collapse. We found that following CS knockdown, the levels of citric acid and isocitric acid decreased compared to the sham group (Figure 4K-L). Additionally, there was a decrease in ATP levels in septic mice after CS knockdown (Figure 4M). TCA cycle metabolites contribute to maintaining mitochondrial homeostasis, and alterations in mitochondrial metabolism could lead to defects in the electron transport chain (ETC). Therefore, we further investigated the impact of CS knockdown on mitochondrial function in septic mice. By measuring the activity of mitochondrial ETC complexes, we observed a decrease in complex activity after CS knockdown (Figure 4N). These findings collectively suggest that CS knockdown exacerbates lung injury and affects mitochondrial function in sepsis mice.

### 5 Inhibition of CS aggravated lung injury and oxidative stress in sepsis mice

Octanoyl-CoA, an inhibitor of CS, was used to observe the changes in lung injury after inhibition of CS. Octanoyl-CoA at 10 mg/mL was the optimal inhibitory dose of CS as determined by quantitative real-time PCR and western blot (Figure S1A-1B). We administered Octanoyl-CoA in sepsis mice. Alveolar wall thickening increased as compared with the sham group (Figure S1C). Octanoyl-CoA administration increased lung injury scores (Figure S1D) and lung wet/dry ratio (Figure S1E). Importantly, Octanoyl-CoA administration significantly increased the survival rate of mice in 72 h (Figure S1F). In addition, the total number of cells and protein concentration levels in BALF were significantly increased after Octanoyl-CoA administration (Figure S1G-1H). Blood gas analysis showed a decrease in PaO_2_ and an increase in PaCO_2_ with Octanoyl-CoA administration (Figure S1I-1J). Octanoyl-CoA administration increased sepsis-induced lung tissues MDA content and MPO activity (Figure S1K-L). In addition, Octanoyl-CoA administration exacerbated the decrease in SOD and GSH activities in sepsis mice (Figure S1M-1N). After Octanoyl-CoA administration, MPO activity was also increased in lung tissues as shown by immunohistochemistry (Figure S1O). This also verified that the inhibition of CS by Octanoyl-CoA administration could aggravate lung injury and oxidative stress in sepsis mice.

### 6 Knockdown of CS aggravated mitochondrial damage in macrophages

We further investigated the effect of SiCS on macrophages after LPS stimulation. Quantitative real-time PCR and western blot showed a decrease in mRNA and protein expression of CS after SiCS treatment (Figure 5A-5B). SiCS inhibited the activity of CS (Figure 5C). Additionally, SiCS reduced the ATP levels and mtDNA copy number contents (Figure 5D-5E). LPS increased the MitoSOX signal pathway, which was further intensified by SiCS treatment (Figure 5F). The protein content of TFAM decreased by SiCS treatment (Figure 5G). Quantitative real-time PCR demonstrated that SiCS increased the levels of Bcl2 while decreasing the levels of BAX and Caspase 9 (Figure 5H-5J). These results indicate that SiCS worsens mitochondrial damage in macrophages and exacerbates cell apoptosis.

**Figure 5.**
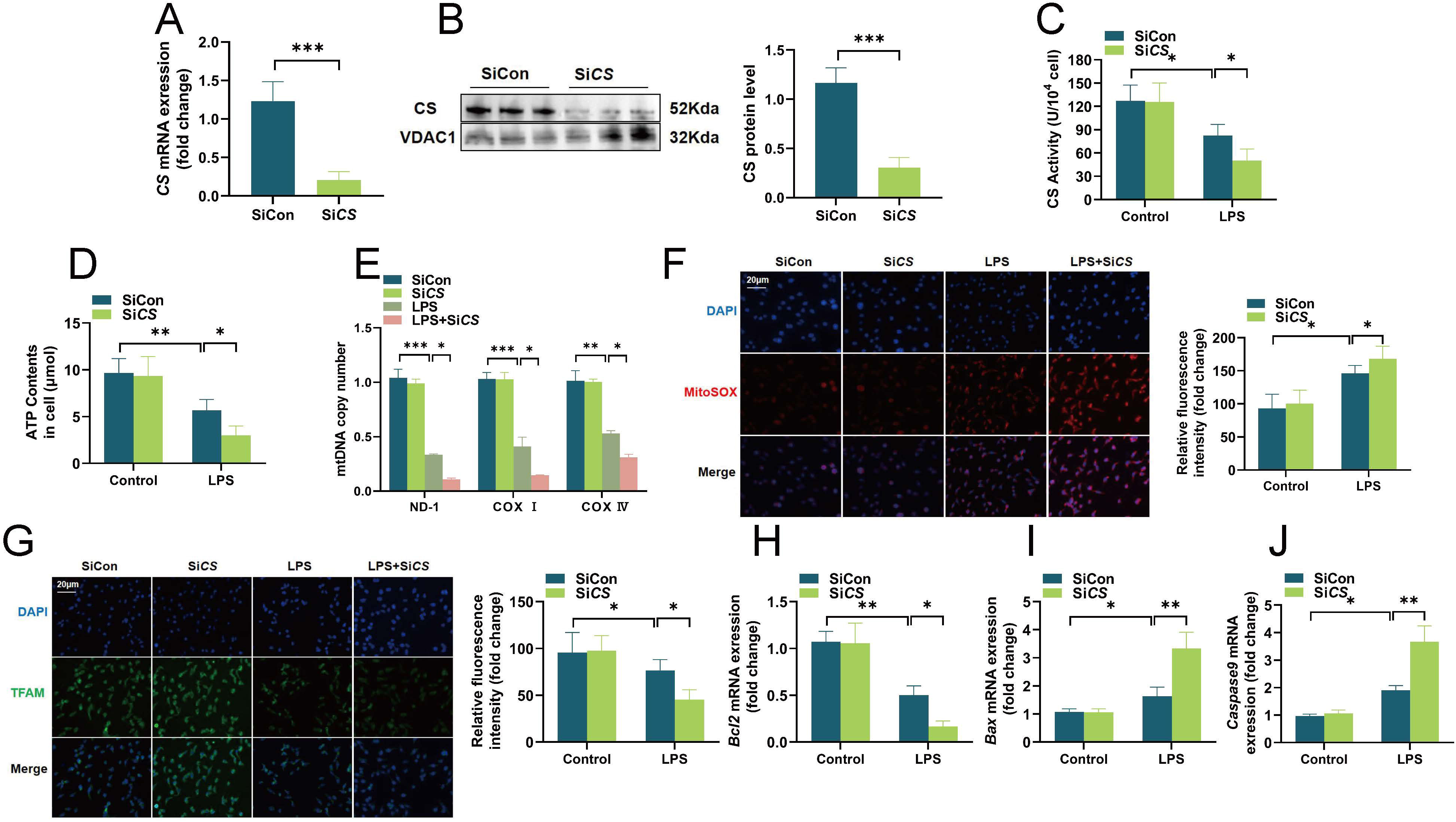
Knockdown of CS aggravated mitochondrial damage in macrophages after LPS stimulation. (A) The mRNA levels of CS in macrophages were detected by quantitative real-time PCR. (B) The protein levels of CS in macrophages were detected by western blot, and the ratio of CS was semi-quantitatively analyzed by ImageJ software. (C) Quantification of CS activity in macrophages. (D) ATP detection kit was used to detect the content of ATP in macrophages. (E) The relative levels of ND-1, COX Ⅰ and COX Ⅳ were used to reflect the mtDNA copy number. (F) MitoSOX Red fluorescent probe (5 µM) was used to detect mitochondrial reactive oxygen species in macrophages, scale bar = 20 µm. (G) Macrophages were stained with the TFAM (green) and DAPI (blue) for immunofluorescence analysis, scale bar = 20 µm. (H-J) The mRNA levels of Bcl2, BAX, and Caspase 9 in macrophages were detected by quantitative real-time PCR. Group comparisons were conducted using analysis of variance (ANOVA). Experiments were repeated at least three times (*n* = 3). \**p* < 0.05, \*\**p* < 0.01, and \*\*\**p* < 0.001.

### 7 Inhibition of CS aggravated mitochondrial damage in macrophages

To further explore the effects of Octanoyl-CoA (the inhibition of CS) in macrophages, we examined the impact of Octanoyl-CoA on macrophages after LPS stimulation. Octanoyl-CoA did not effect on the vitality of macrophages (Figure S2A). It was observed that the damage became severe 6 h after LPS stimulation (Figure S2B). By measuring ATP levels and CS activity, it was found that 5mM Octanoyl-CoA began to inhibit CS (Figure S2C-2D). Quantitative real-time PCR and western blot showed that Octanoyl-CoA inhibited the mRNA and protein expression of CS (Figure S2E-2F). Octanoyl-CoA also inhibited the activity of CS (Figure S2G). Octanoyl-CoA reduced the ATP levels (Figure S2H). Transmission electron microscopy revealed that Octanoyl-CoA exacerbated mitochondrial damage (Figure S2I). LPS led to an increase in the MitoSOX signal pathway, which was further compounded by Octanoyl-CoA treatment (Figure S2J). Flow cytometry also indicated that Octanoyl-CoA significantly increased LPS-induced apoptosis in macrophages (Figure S2K). Western blot showed that Octanoyl-CoA increased the expression of Bcl2 while decreasing the expression of BAX (Figure S2L). Quantitative real-time PCR demonstrated that Octanoyl-CoA increased the levels of Bcl2 while decreasing the levels of BAX and Caspase 9 (Figure S2M). The above results indicate that after inhibiting CS, Octanoyl-CoA exacerbated mitochondrial damage in macrophages and intensified cell apoptosis.

### 8 Overexpression of CS alleviated mitochondrial damage in macrophages

To further elucidate the impact of CS in macrophages after LPS stimulation, we investigated whether CS overexpression (CS-OE) could alleviate the effects in macrophages after LPS stimulation. Quantitative real-time PCR and western blot revealed increased mRNA and protein expression of CS after CS-OE (Figure 6A-6B). CS-OE enhanced CS activity (Figure 6C). Quantitative real-time PCR confirmed that CS-OE significantly improved the RNA levels of nuclear-encoded mitochondrial genes and mtDNA-encoded genes of the ETC complexes (Figure 6D-6F). LPS led to an increase in the MitoSOX signal, which was attenuated by CS-OE treatment (Figure 6G). And the protein content of COX I increased after CS-OE treatment (Figure 6H). Additionally, CS-OE increased the mtDNA copy number content (Figure 6I). These results indicate that CS-OE can alleviate mitochondrial damage and reduce cellular injury.

**Figure 6.**
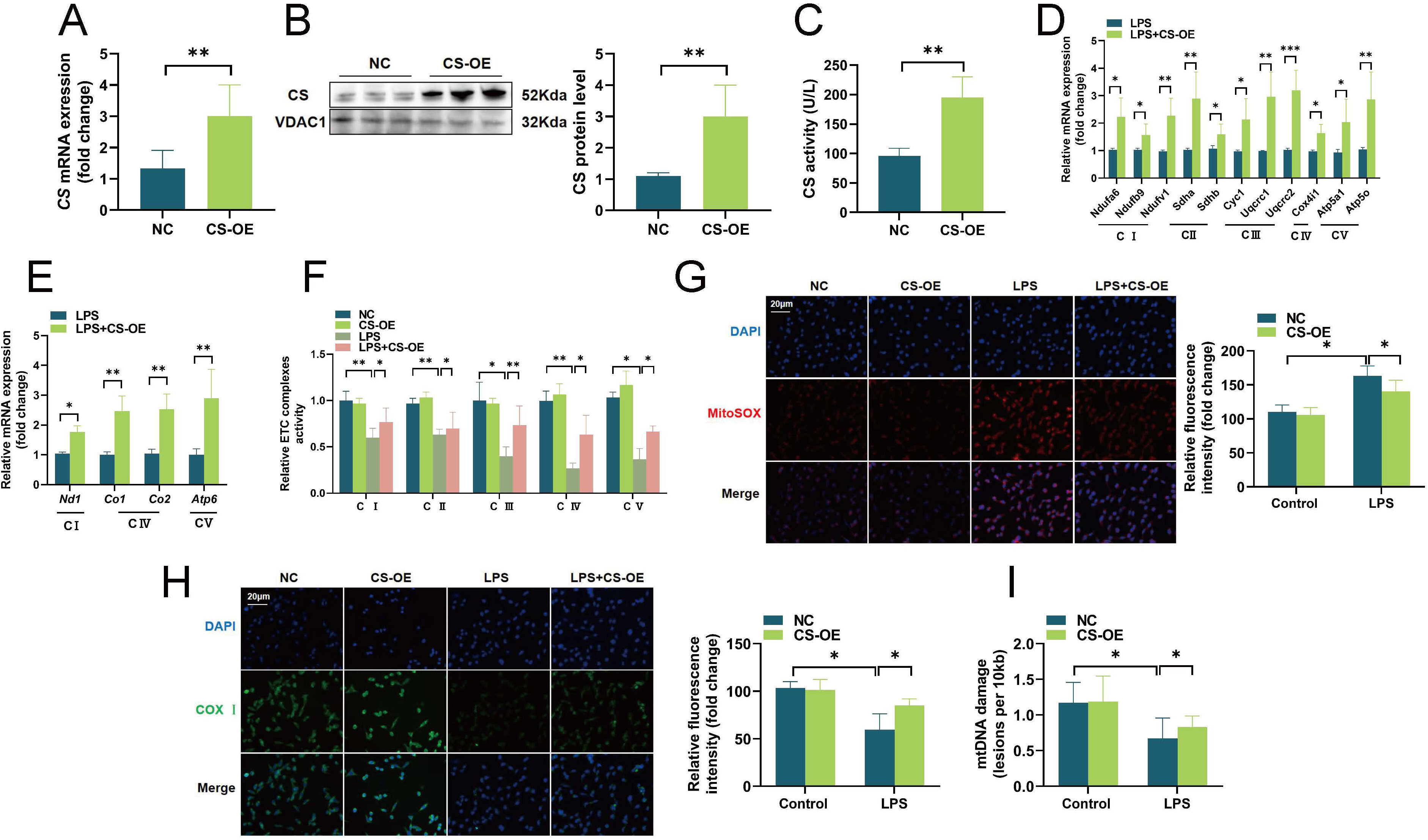
Overexpression of CS alleviated mitochondrial damage in macrophages after LPS stimulation. (A) The mRNA levels of CS in macrophages were detected by quantitative real-time PCR. (B) The protein levels of CS in macrophages were detected by western blot, and the ratio of CS was semi-quantitatively analyzed by ImageJ software. (C) Quantification of CS activity in macrophages. (D-E) The mRNA levels of the mitochondrial complex in macrophages were detected by quantitative real-time PCR. (F) The relative levels of mitochondrial complex activity. (G) MitoSOX Red fluorescent probe (5 µM) was used to detect mitochondrial reactive oxygen species in macrophages, scale bar = 20 µm. (H) Macrophages were stained with the COX Ⅰ (green) and DAPI (blue) for immunofluorescence analysis, scale bar = 20 µm. (I) mtDNA damage detection in macrophages after LPS stimulation. Group comparisons were conducted using analysis of variance (ANOVA). Experiments were repeated at least three times (*n* = 3). \**p* < 0.05, \*\**p* < 0.01, and \*\*\**p* < 0.001.

### 9 CS regulated mitochondrial metabolism through the TCA cycle in macrophages

To further understand the relationship between CS and the pathogenesis of sepsis, we conducted metabolomic analysis on mitochondria. Bioinformatic analysis of differentially expressed genes revealed that upregulated genes were enriched in metabolic pathways including the TCA cycle (Figure 7A-7B). We performed integrated analysis of the transcriptome (differentially expressed genes with Benjamini-Hochberg corrected p < 0.05 and |log2[fold change] | ≥ 1) and metabolites with significantly variable projections in orthogonal partial least squares discriminant analysis > 1.2). In sepsis patients, the downstream metabolites of CS, including citric acid and isocitric acid, showed decreased abundance (Figure 7C-7D). In macrophages after LPS stimulation, after SiCS, citric acid and isocitric acid decreased more severely (Figure 7E-7F).

**Figure 7.**
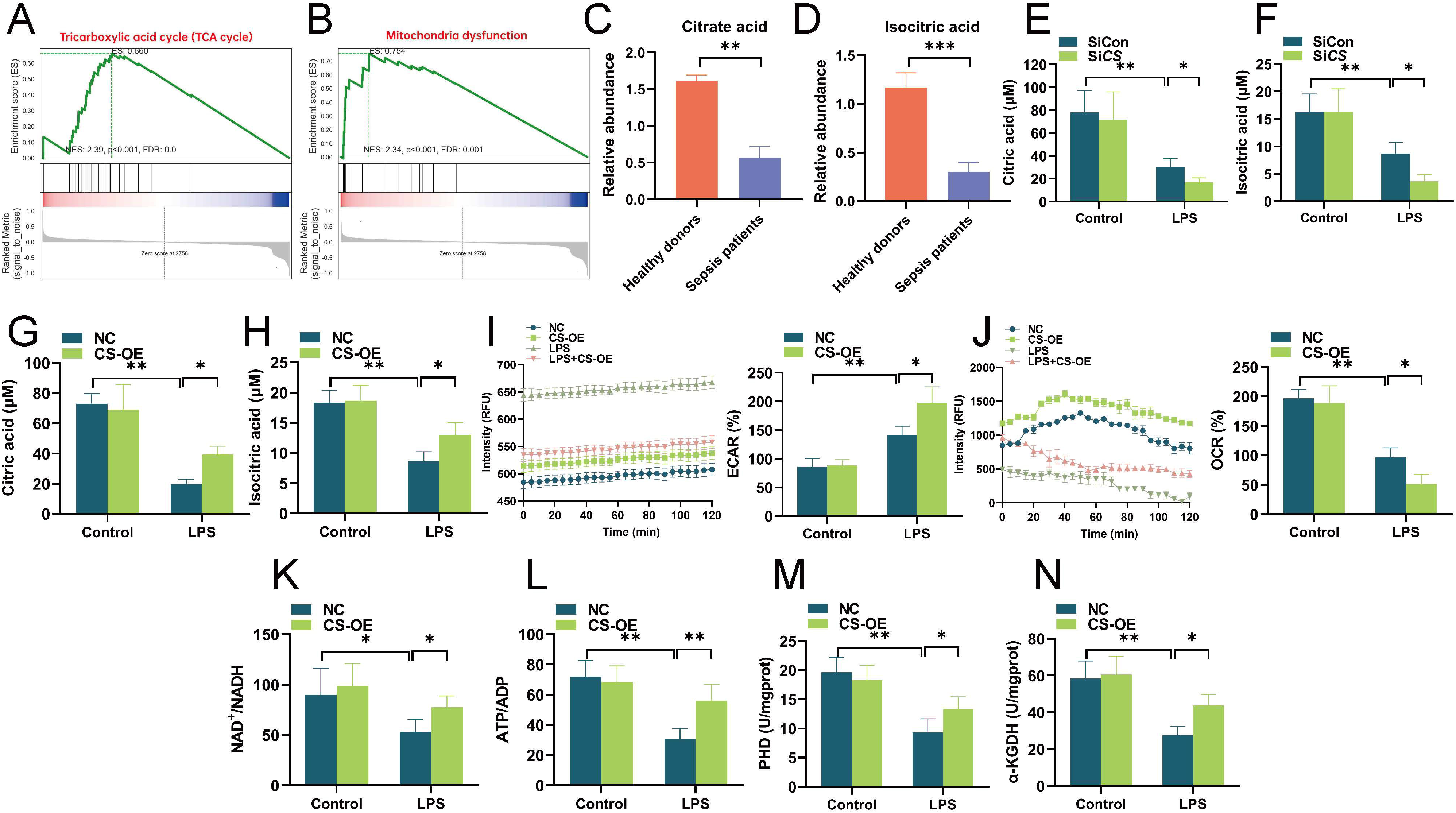
CS regulated mitochondrial metabolism through the TCA cycle in macrophages. (A-B) A representative GSEA enrichment plot of the transcription profile that was used to identify the CS-activated signaling pathway. (C-D) Relative abundances of citric acid and isocitric acid in sepsis patients. (E-F) Relative abundances of citric acid and isocitric acid in macrophages after LPS stimulation. (G-H) Relative abundances of citric acid and isocitric acid in macrophages after CS-OE. (I-J) The ECAR and OCR were measured in macrophages after CS-OE. (K-N) NAD^+^/NADH, ATP/ADP, PHD, and α-KGDH activities in macrophages. Group comparisons were conducted using analysis of variance (ANOVA). Experiments were repeated at least three times (*n* = 3). \**p* < 0.05, \*\**p* < 0.01, and \*\*\**p* < 0.001.

TCA cycle metabolites contribute to the maintenance of mitochondrial homeostasis, and alterations in mitochondrial metabolism can lead to impaired mitochondrial respiration, mtDNA damage, and ETC defects. Therefore, we further investigated the effect of CS-OE on mitochondrial function in macrophages. Citric acid and isocitric acid were increased in macrophages after CS-OE (Figure 7G-7H). We also found that ECAR decreased, and OCR increased after CS-OE (Figure 7I-7J). Moreover, it also increased the NAD^+^/NADH, ATP/ADP, PHD, and α-KGDH activities after CS-OE (Figure 7K-7N). Collectively, these data indicated that CS regulated mitochondrial homeostasis in macrophages.

### 10 Supplementation with the CS downstream metabolite Citric acid attenuated macrophage injury

In the TCA cycle, CS catalyzes the condensation of acetyl-Coa with oxaloacetate to yield citrate. Therefore, we wondered whether the impaired TCA cycle in macrophages could be reactivated by supplementation of Citric acid, a key metabolic downstream product. To test this hypothesis, we found that LPS resulted in attenuated Mito tracker signaling, whereas Citric acid increased Mito tracker signaling (Figure 8A). We further explored the role of Citric acid in the coordination of glycolysis and mitochondrial respiration by measuring ECAR and OCR. Citric acid resulted in decreased levels of glycolysis and enhanced mitochondrial respiration (Figure 8B-8C). NAD^+^/NADH, ATP/ADP, PHD, and α-KGDH activities were significantly upregulated after Citric acid administration (Figure 8D-8G). In addition, Citric acid increased mitochondrial complex protein content (Figure 8H). Citric acid treatment increaesed the mRNA of Bcl2 while decreasing BAX and Caspase 9 (Figure 8I). These results suggest that supplementation with Citric acid, a downstream metabolite of CS, prevents macrophage damage.

**Figure 8.**
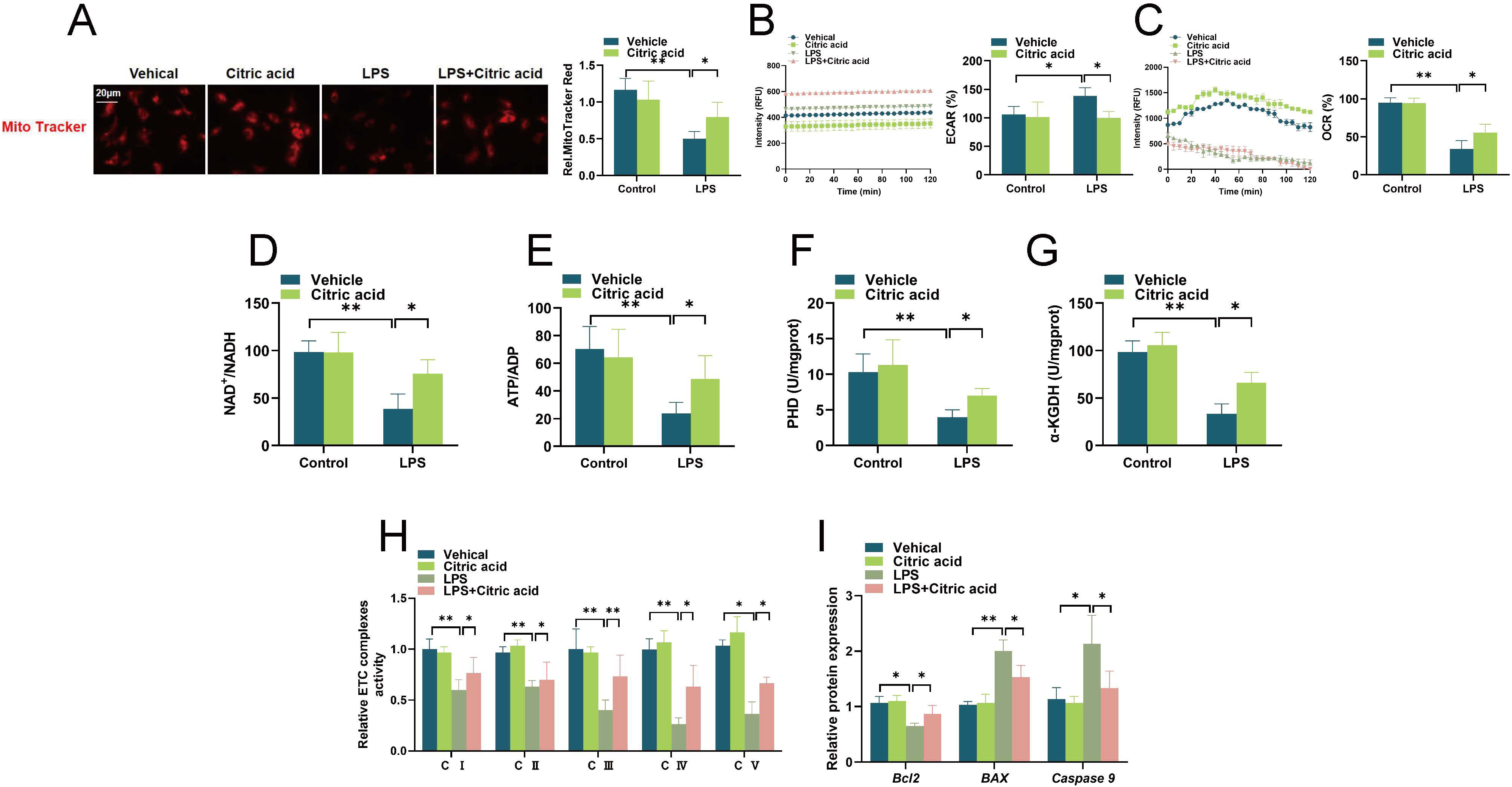
Supplementation with the CS downstream metabolite Citric acid attenuated macrophage injury. (A) Mito tracker was used to detect macrophages, scale bar = 20 µm. (B-C) The ECAR and OCR were measured in macrophages after LPS stimulation. (D-G) NAD^+^/NADH, ATP/ADP, PHD, and α-KGDH activities in macrophages. (H) The relative levels of mitochondrial complex activity. (I) The mRNA levels of Bcl2, BAX, and Caspase 9 in macrophages were detected by quantitative real-time PCR. Group comparisons were conducted using analysis of variance (ANOVA). Experiments were repeated at least three times (*n* = 3). \**p* < 0.05 and \*\**p* < 0.01.

### 11 Supplementation with Citric acid attenuated macrophage injury after SiCS

We further tested whether Citric acid supplementation attenuated macrophage damage after SiCS. The protein content of TFAM increased after Citric acid treatment (Figure 9A). We found that SiCS resulted in attenuated Mito tracker signaling, whereas Citric acid increased Mito tracker signaling (Figure 9B). Additionally, SiCS decreased the ATP levels and mitochondrial complex activity, and Citric acid increased the ATP levels and mitochondrial complex activity (Figure 9C-9D). SiCS decreased the mtDNA copy number content and Citric acid increased the mtDNA copy number content (Figure 9E). SiCS decreased the activities of NAD^+^/NADH, ATP/ADP, PHD, and α-KGDH and they were significantly upregulated after citric acid administration (Figure 9F-9I).

**Figure 9.**
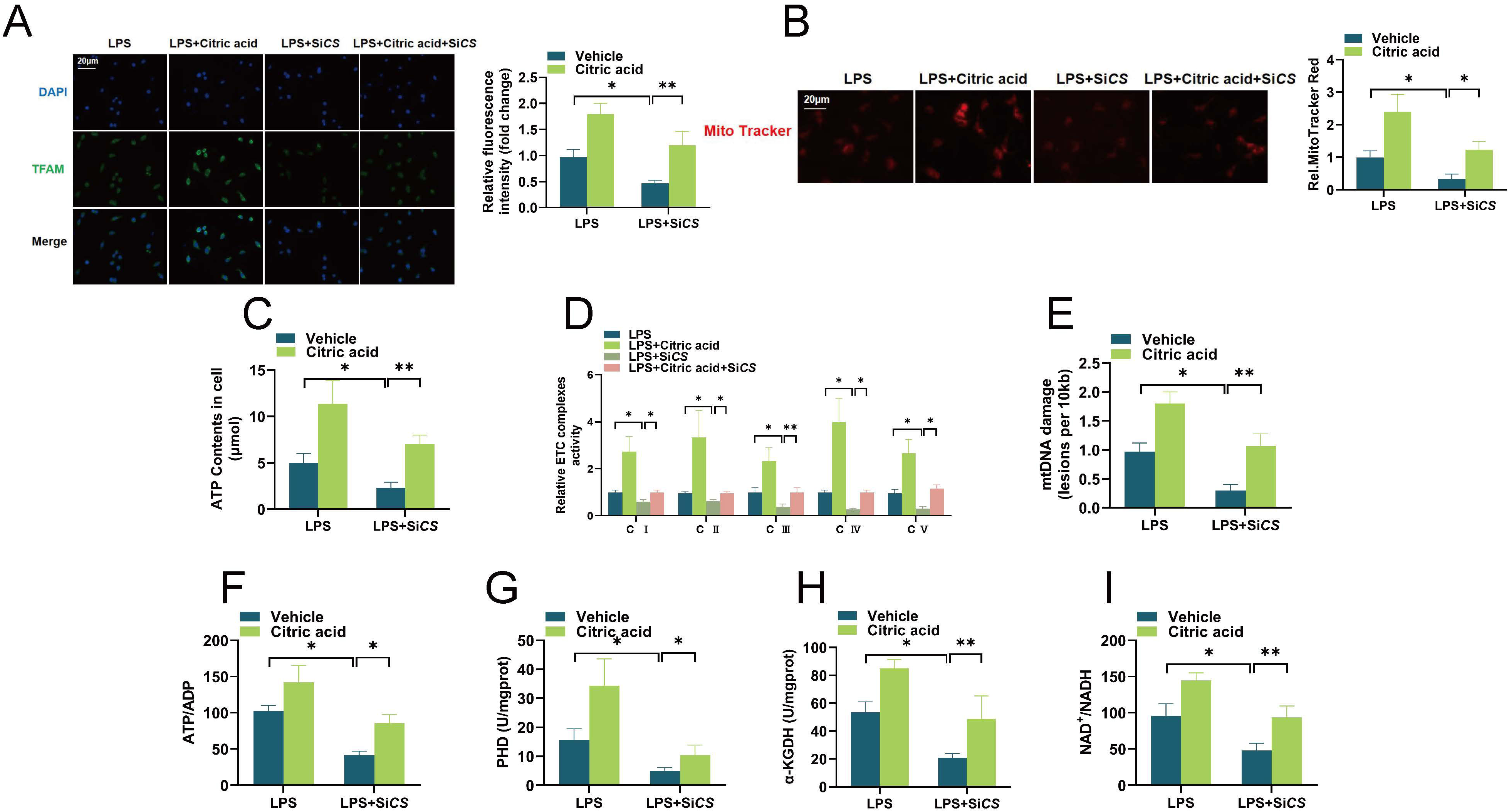
Supplementation with Citric acid attenuated macrophage injury after SiCS. (A) Macrophages were stained with the TFAM (green) and DAPI (blue) for immunofluorescence analysis, scale bar = 20 µm. (B) Mito tracker was used to detect macrophages, scale bar = 20 µm. (C) ATP detection kit was used to detect the content of ATP in macrophages. (D) The relative levels of mitochondrial complex activity. (E) mtDNA damage detection in macrophages. (F-I) NAD^+^/NADH, ATP/ADP, PHD, and α-KGDH activities. Group comparisons were conducted using analysis of variance (ANOVA). Experiments were repeated at least three times (*n* = 3). \**p* < 0.05, \*\**p* < 0.01, and \*\*\**p* < 0.001.

### 12 Supplementation with Citric acid attenuated lung injury after inhibiting CS in sepsis mice

We administered Citric acid and Octanoyl-CoA in sepsis mice. Citric acid reduced the Octanoyl-CoA-induced increase in alveolar wall thickening compared with the Sham group (Figure 10A). Citric acid reduced Octanoyl-CoA -induced lung injury score (Figure 10B) and lung wet/dry ratio increase (Figure 10C). In addition, Citric acid reduced the significant increase in total cell numbers and protein concentration levels in BALF induced by Octanoyl-CoA (Figure 10D-10E). Blood gas analysis showed that Citric acid reduced the Octanoyl-CoA-induced decrease in PaO_2_ and increase in PaCO_2_ (Figure 10F-10G). Additionally, Citric acid reduced the ATP levels (Figure 10H).

**Figure 10.**
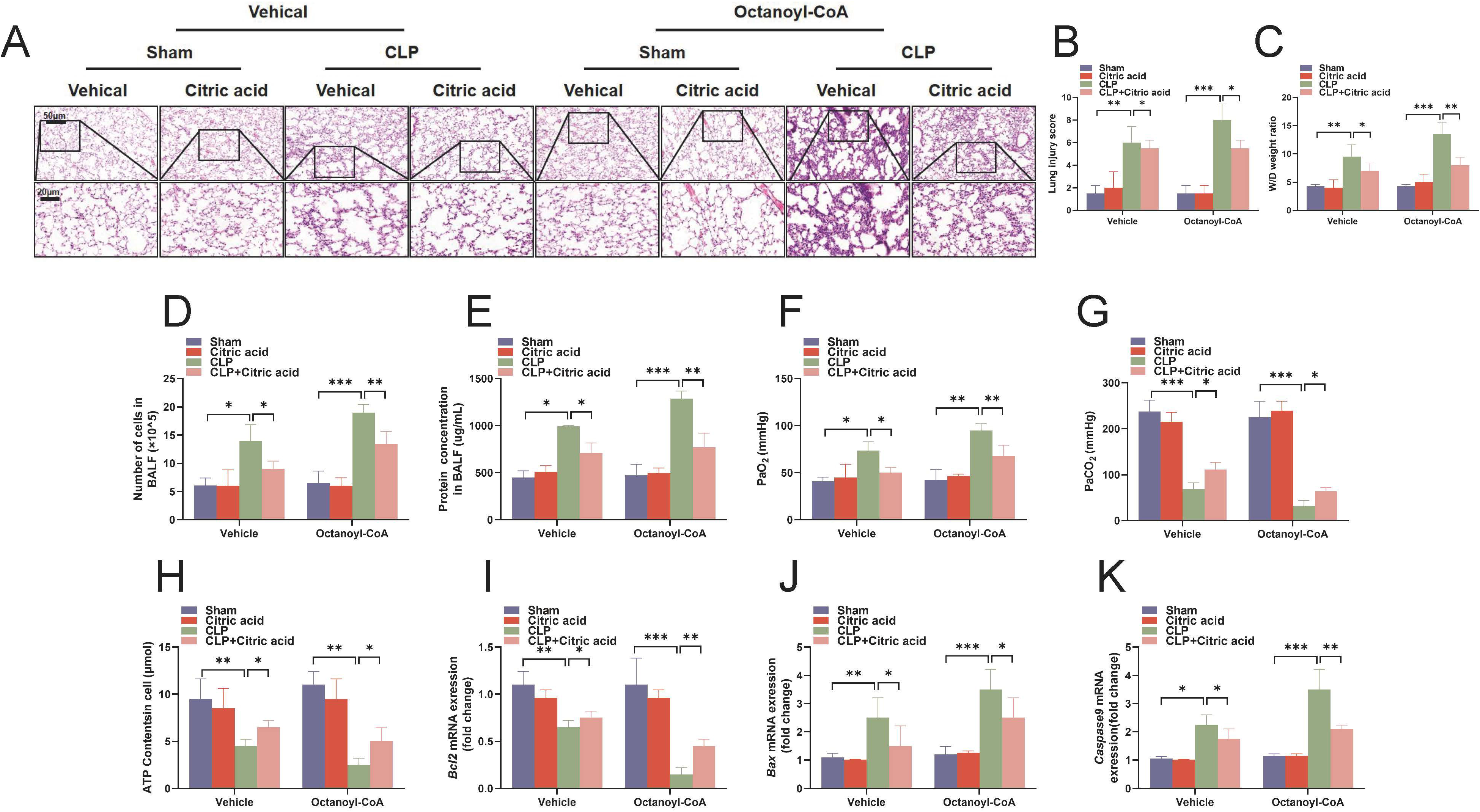
Supplementation with Citric acid attenuated lung injury after inhibiting CS in sepsis mice. (A) Hematoxylin and eosin staining were performed to evaluate the lung histopathological changes (magnification, × 200). (B) Mouse lung injury score was calculated. (C) The wet-to-dry ratio of lung tissues. (D) The total number of cells in BALF. (E) The total protein concentrations of BALF. (F-G) Blood gas analysis of sepsis mice. (H) ATP detection kit was used to detect the content of ATP in sepsis mice. (I-K) The mRNA levels of Bcl2, BAX, and Caspase 9 in sepsis mice were detected by quantitative real-time PCR. Group comparisons were conducted using analysis of variance (ANOVA). Experiments were repeated at least three times (*n* = 6). \**p* < 0.05, \*\**p* < 0.01, and \*\*\**p* < 0.001.

Quantitative real-time PCR demonstrated that Citric acid increased the levels of Bcl2 while decreasing the levels of BAX and Caspase 9 (Figure 10I-10K). This data provides compelling evidence that supplementation of CS downstream metabolite Citric acid prevents lung injury induced by sepsis.

## Discussion

Sepsis is an extremely complex cascade of events, involving fluid and cellular responses, inflammatory and anti-inflammatory processes, and circulatory abnormalities (15). Over time, bacterial resistance to antibiotics continues to increase, making treating sepsis increasingly challenging. Sepsis may present with atypical symptoms in its early stages, often being overlooked, thereby complicating the early diagnosis of sepsis and delaying treatment initiation (1). Sepsis can lead to severe systemic inflammatory responses, resulting in multiple organ dysfunction, further complicating treatment. The severity of sepsis caused by different pathogens varies, with some pathogens showing poor responses to treatment. Individuals using immunosuppressants or those with immune deficiencies may exhibit a poorer response to sepsis treatment. The treatment of sepsis may require a comprehensive approach involving antibiotics, hemodynamic support, immunomodulation, and other methods, making the treatment regimen complex (16, 17). Sepsis treatment can lead to sequelae and complications, such as long-term organ dysfunction and changes in mental status in patients. Therefore, it is crucial to delve deeper into the pathogenesis of sepsis and identify new targets for treatment.

Mitochondria are specialized intracellular structures surrounded by a double membrane, serving as the hub of cellular metabolism and participating in a series of crucial cellular processes, including oxidative phosphorylation, the tricarboxylic acid cycle, maintaining intracellular calcium homeostasis, as well as the generation and maintenance of reactive oxygen species (ROS) (18, 19). During the development of sepsis, mitochondria undergo morphological and functional damage in multiple ways. Liang et al. found irregular mitochondrial morphology in cardiomyocytes of a septic animal model, characterized by significant swelling and vacuolar degeneration, with evident mitochondrial cristae rupture observed in the swollen mitochondria (20). Similar results were reported by Zhang et al (21). In addition to abnormal mitochondrial morphology, sepsis also impairs mitochondrial function, leading to a significant decrease in mitochondrial membrane potential. However, the specific mechanisms still require further investigation.

The tricarboxylic acid cycle (TCA cycle), also known as the citric acid cycle or Krebs cycle, is one of the most important metabolic pathways in living organisms, primarily occurring in the matrix of mitochondria. The TCA cycle is the process of converting carbon sources (such as glucose, fatty acids) into energy, generating ATP and providing coenzymes like NADH and FADH₂ needed by cells (22, 23). There is a close relationship between the TCA cycle and sepsis, mainly involving the regulation of cellular metabolism and the balance of energy production. In a state of sepsis, cellular metabolism is disrupted, leading to impaired function of the TCA cycle. Under normal circumstances, the TCA cycle converts metabolic products into NADH and FADH₂, providing the electrons needed for ATP production within the cell (24).

However, in sepsis, due to increased inflammatory responses and oxidative stress, the TCA cycle is disrupted, causing disturbances in cellular energy metabolism. Oxidative stress within mitochondria during sepsis may damage mitochondrial DNA and membrane structures, affecting the function of the TCA cycle, thereby impacting cell survival and function (25). Additionally, Peace et al. found in a mouse model of sepsis that the TCA cycle product OI has been shown to improve survival rates and prevent LPS-induced lethality (26). Jorge et al. found that beta-glucans also increase the expression of succinate dehydrogenase (SDH), contributing to the integrity of the TCA cycle, enhancing the innate immune response after secondary stimulation, making it an attractive therapeutic target in sepsis (27).

CS is one of the key enzymes involved in the TCA cycle. This enzyme plays a crucial role in the initial stages of the TCA cycle by catalyzing the formation of citrate from acetyl coenzyme A and oxaloacetate, marking the beginning of the TCA cycle (28). Therefore, Citrate synthase plays a vital role in the TCA cycle by catalyzing the synthesis of citrate, initiating and regulating the entire TCA cycle process to provide the cell with the necessary energy and metabolites. Our research has revealed a novel mechanism underlying the occurrence of sepsis, targeting CS to reconstruct macrophage mitochondrial TCA cycle, increase cellular energy generation, and improve sepsis-induced lung injury. In sepsis patients, CS is expressed at low levels and is positively correlated with lung function indicators. The AUC for distinguishing sepsis patients from healthy donors is 0.9727. Knocking down CS or inhibiting its expression in septic mice worsened lung injury and oxidative stress. In macrophages, inhibiting CS expression affected the TCA cycle, exacerbating cell apoptosis, while overexpressing CS promoted the TCA cycle, alleviated cell apoptosis, enhanced cellular energy generation, and reduced oxidative stress levels. Supplementation of citric acid (a downstream metabolite of CS) helped alleviate mitochondrial damage, promote the TCA cycle, and supplementing citric acid could alleviate lung injury in septic mice.

This research outcome paves the way for new avenues in the study and therapeutic intervention of sepsis. Targeting CS and macrophage tricarboxylic acid cycle represent a novel approach to regulating cellular metabolism and immune responses in the context of sepsis. Further exploration of the molecular mechanisms of CS in the pathogenesis of sepsis may lay the groundwork for developing targeted therapies aimed at enhancing cellular energetics and improving clinical outcomes in sepsis patients. Additionally, citric acid, as a downstream metabolite of CS, holds great promise in improving mitochondrial damage and promoting TCA cycle function. Supplementation of citric acid in a mouse model can alleviate sepsis-induced lung injury, highlighting the therapeutic efficacy of targeting the metabolic pathways mediated by CS in sepsis management. These results underscore the potential of citric acid as a therapeutic agent in alleviating sepsis-related complications.

## Authors’ contributions

Sun contributed equally to this article. Dr. Wang conceived and designed these experiments. Jin collected clinical data. Sun performed the main experiments. Jin analyzed the bioinformatics data. Wang organized the clinical data. Dr. Wang revised the article. All authors read and approved of the final manuscript.

## Informed Consent Statement

Not applicable.

## Data Availability Statement

The original data are available from the corresponding author upon reasonable request.

## Supplementary Figure legends

**Figure S1** Inhibition of CS aggravated lung injury and oxidative stress in sepsis mice. (A) The mRNA levels of CS in lung tissues were detected by quantitative real-time PCR. (B) The protein levels of CS in mice lung tissues were detected by western blot, and the ratio of CS was semi-quantitatively analyzed by ImageJ software. (C) Hematoxylin and eosin staining were performed to evaluate the lung histopathological changes (magnification, × 200). (D) Mouse lung injury score was calculated. (E) The wet-to-dry ratio of lung tissues. (F) The accumulative survival curves of sepsis mice (*n* = 10). (G) The total number of cells in BALF. (H) The total protein concentrations of BALF. (I-J) Blood gas analysis of sepsis mice. (K-N) The levels of MDA, MPO, SOD, and GSH in lung tissues were measured by the appropriate detection kit. (O) CS immunohistochemistry and semi-quantitative analysis of CS immunohistochemistry (magnification, ×200). Group comparisons were conducted using analysis of variance (ANOVA). Experiments were repeated at least three times (*n* = 6). \**p* < 0.05, \*\**p* < 0.01, \*\*\**p* < 0.001, and \*\*\*\**p* < 0.001.

**Figure S2** Inhibition of CS aggravated mitochondrial damage in macrophages after LPS stimulation. (A) CCK8 assay was performed to detect the toxicity of CS to macrophages. (B) CCK8 assay was performed to detect the toxicity of LPS to macrophages. (C-D) The optimal dose of CS for macrophages was detected by the CCK8 method. (E) The mRNA levels of CS in macrophages were detected by quantitative real-time PCR. (F) The protein levels of CS in macrophages were detected by western blot, and the ratio of CS was semi-quantitatively analyzed by ImageJ software. (G) Quantification of CS activity in macrophages. (H) ATP detection kit was used to detect the content of ATP in macrophages. (I) Electron microscopic analysis of mitochondrial structure in macrophages. (J) MitoSOX Red fluorescent probe (5 µM) was used to detect mitochondrial reactive oxygen species in macrophages, scale bar = 20 µm. (K) The apoptosis of macrophages was measured by flow cytometry. (L) The protein levels of Bcl2 and BAX in macrophages were detected by western blot, and the ratio of Bcl2 and BAX was semi-quantitatively analyzed by ImageJ software. (M) The mRNA levels of pro-apoptotic and anti-apoptotic genes in macrophages were detected by quantitative real-time PCR. Group comparisons were conducted using analysis of variance (ANOVA). Experiments were repeated at least three times (*n* = 3). \**p* < 0.05, \*\**p* < 0.01, \*\*\**p* < 0.001, and \*\*\*\**p* < 0.001. ns means no significant difference.

