## Supplementary material for "Citrate synthase improves sepsis-induced lung injury by reconstructing the mitochondrial tricarboxylic acid cycle of macrophages": Table S1

**Table S1 The characteristics of heathy donors and sepsis patients**

|  | **Sepsis patients**  **(*n* = 76)** | **healthy donors**  **(*n* = 89)** | **Inspection value**  **(t, χ2 or z)** | ***p* value** |
| --- | --- | --- | --- | --- |
| **Age (range)** | 75.4 ± 5.5  (67-88) | 74.1 ± 8.6  (55-91) | 1.456 | 0.105 |
| **Sex (%)** | Male = 46 (60.52%) | Male = 54 (60.67%) | 0.217 | 0.626 |
| **Physiological parameters**  **during exposure window**  PCT [ng/ml; IQR]  CRP (mg/L; range)  Serum lactate concentration [mg/dL; IQR]  MAP [mmHg; IQR]  PaO2/FiO2 (mmHg)  Creatinine [μmol/L; IQR] | 11.8 [2.45, 17.53]  103.1 ± 89.9 (6.8 to 354.9)  1.5 [1.16, 2.67]  84.12 [67.57, 90.13]  122.32 ± 61.03 (55.5 to 238.1)  128.41 [87.56, 266.17] | 1.2 [0.52, 3.54]  37.5 ± 21.5 (6.7 to 124.3)  1.4 [0.94, 3.70]  83.00 [74.67, 97.54]  232.35 ± 154.87 (127.1 to 498.2)  145.70 [66.48, 244.63] | -1.067  1.637  -0.545  -0.377  3.854  -1.467 | 0.017  0.007  0.654  0.864  0.003  0.578 |
| **APACHE II score (range)** | 27.6 ± 6.8 (8 to 41) | 14.5 ± 4.0 (8 to 23) | 7.668 | <0.001 |
| **SOFA score (range)** | 13.6 ± 4.0 (4 to 20) | 5.5 ± 2.4 (1 to 9) | 7.467 | <0.001 |
| **ICU length of stay (d;IQR)** | 14 [8, 21] | 20.5 [14.5, 34.75] | -1.345 | 0.543 |
| **Death within 28 days (%)** | 24 (30.4%) | 1(0.9%) | 3.577 | 0.355 |

Variables were presented as mean ± SD, number of patients (n) or median [IQR]. PCT, Procalcitonin; CRP, C-reactive protein: MAP, mean arterial pressure; APACHE II, Acute Physiology and Chronic Health Evaluation II; SOFA, Sequential Organ Failure Assessment; IQR, interquartile range.
