## Supplementary material for "Citrate synthase improves sepsis-induced lung injury by reconstructing the mitochondrial tricarboxylic acid cycle of macrophages": Table S2

**Table S2. Primer sequences of the target genes**

| **Target Gene** | **Primer** | **Primer Sequences (5’ to 3’)** |
| --- | --- | --- |
| GAPDH (mouse) | Forward  Reverse | AGGTCGGTGTGAACGGATTTG  TGTAGACCATGTAGTTGAGGTCA |
| CS (mouse) | Forward  Reverse | CTTTGGTAATATGCTGGG  ATGTTGCCTATACGTTGTG |
| NDUFB9 (mouse) | Forward  Reverse | GTGGTGCGTCCAGAGAGAC  GGCCTTCGCCATATCCTTTTC |
| NDUFV1 (mouse) | Forward  Reverse | AGGATGAAGACCGGATTTTCAC  CAGTCACCTCGACTCAGGGA |
| SDHA (mouse) | Forward  Reverse | CAAACAGGAACCCGAGGTTTT  CAGCTTGGTAACACATGCTGTAT |
| SDHB (mouse) | Forward  Reverse | ACAGCTCCCCGTATCAAGAAA  GCATGATCTTCGGAAGGTCAA |
| CYC1 (mouse) | Forward  Reverse | CTTCGCGGGGTAGTGTTGG  GGCCAGACTTCGACGACAA |
| Cox4i1 (mouse) | Forward  Reverse | TTGGCAAGAGAGCCATTTCT  GCGTAAGTGGGGAAAGCATA |
| Atp5a1 (mouse) | Forward  Reverse | GCCCTCGGTAATGCTATTGA  GCCCTCGGTAATGCTATTGA |
| mt-Nd1 (mouse) | Forward  Reverse | TCCGAGCATCTTATCCACGC  GTATGGTGGTACTCCCGCTG |
| mt-Cox1(mouse) | Forward  Reverse | CTACCCACCTCTAGCCGGAA  TGTTATGGCTGGGGGTTTCA |
| mt-Cox2 (mouse) | Forward  Reverse | ACCGAGTCGTTCTGCCAATA  ATTTAGTCGGCCTGGGATGG |
| mt-Atp6 (mouse) | Forward  Reverse | ACGCCTAATCAACAACCGTC  TTCGTCCTTTTGGTGTGTGGA |
| Bcl2 (mouse) | Forward  Reverse | TTGAGCATAGTGGTTGGAC  TGGAGGAGGAGAGGAGAA |
| BAX (mouse) | Forward  Reverse | ACGCTGTCCAAACCGTC  TTAACCAGGTCAAGAGCAC |
| Caspase 9 (mouse) | Forward  Reverse | CTCTTGAAATAGCAGACGGG  CAGGTGATAACTAAAACGGTCA |
