## Supplementary figures and images for "Citrate synthase improves sepsis-induced lung injury by reconstructing the mitochondrial tricarboxylic acid cycle of macrophages"

### Figure S1

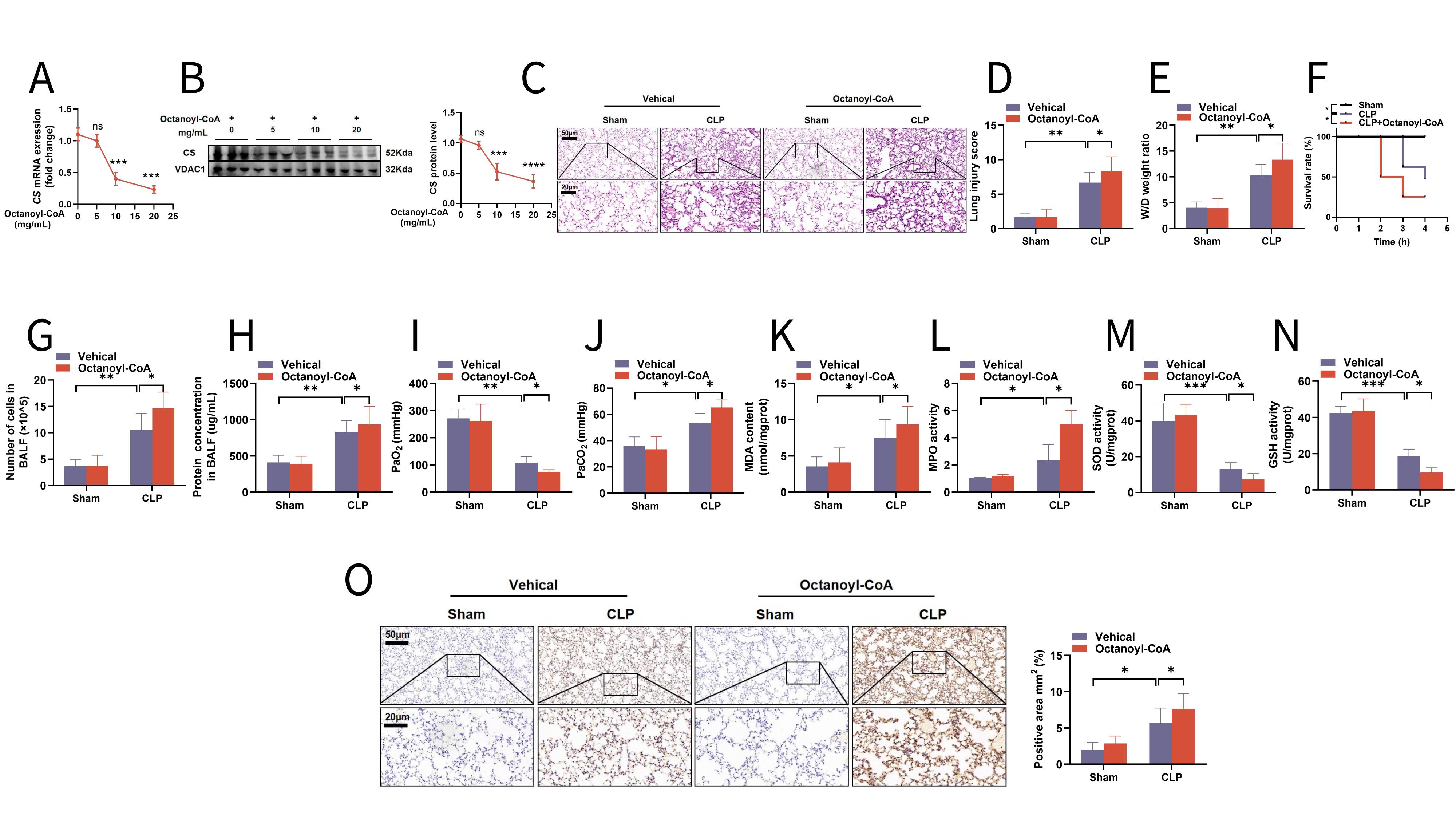

### Figure S2

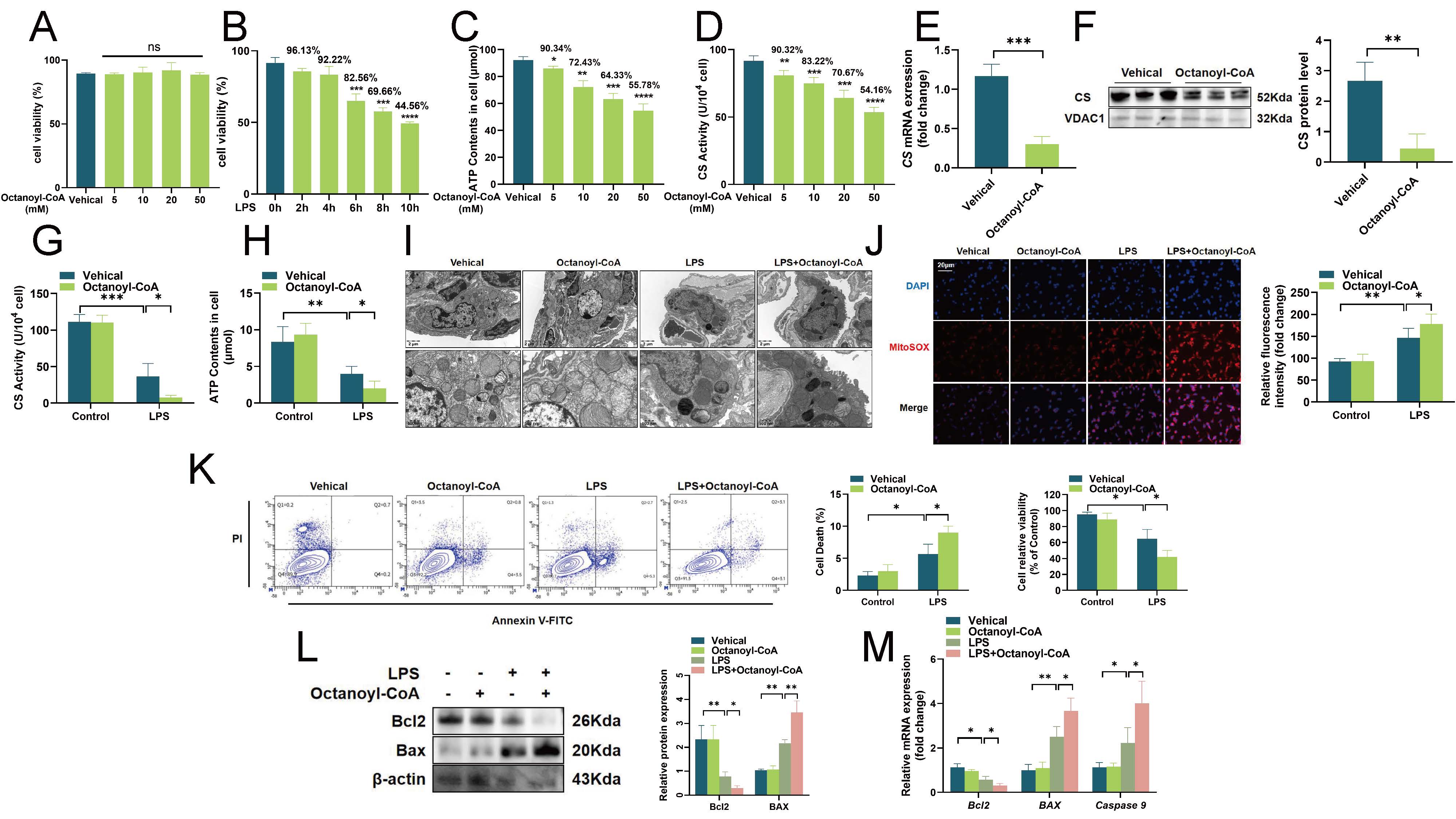
